# Transcriptomic assessment of the Black Judaicus scorpion (*Hottentotta judaicus*) toxin arsenal and its potential bioeconomic value

**DOI:** 10.1101/2025.07.09.664027

**Authors:** Tim Lüddecke, Josephine Dresler, Sabine Hurka, Yachen Wang, Tamara Pohler, Yuri Simone, Jonas Krämer, Andreas Vilcinskas, Volker Herzig

## Abstract

Scorpion venoms are likewise a medical burden as well as a source of novel bioresources. Despite their important dual role, the venoms of most scorpion species remain under- or unstudied. Among these is the venom of the Black Judaicus scorpion, *Hottentotta judaicus* (Simon, 1872), a common yet neglected species native to the Middle East. Here, we employ venom gland transcriptomics to investigate its toxin-encoding precursor profile to gain insight into its toxin repertoire. The venom was found to be composed primarily of various short scorpion toxins, long scorpion toxins from the 3 C-C as well as the 4 C-C type, and enzymatic components. Minor components include, defensins and putative antimicrobial peptides. Several identified toxins show similarity to known neurotoxins from lethal buthids or to toxins with translational value in biomedicine, agriculture, and industrial production, thus rendering *H. judaicus* both, a potential health concern and source of novel bioresources. Our work provides an extended perspective on the venom profile of this species and represents a basis for future follow-up studies.

## 1. Introduction

Scorpions are an ancient lineage of predatory arachnids that employ venom mainly to subdue their prey and deter predators (Es-Saadi et al., 2025; Lüddecke et al., 2022; Simone and van der Meijden, 2021). Venoms are chemically complex and often contain hundreds of potent and highly selective biomolecules referred to as toxins (Casewell et al., 2013; Fry et al., 2009; Hayes et al., 2025; Herzig, 2019; Nelsen et al., 2014). The toxin repertoire of scorpions is primarily composed of polypeptides (proteins and peptides) that often target receptors and ion channels involved in neurological signal transduction and hence cause neurotoxic symptoms (Xia et al., 2023).

Scorpions have a dualistic role for humanity. On one hand, they are an important global health concern (Chippaux and Goyffon, 2008; Isbister and Bawaskar, 2014; Kazemi et al., 2024; Pucca et al., 2025). Their painful sting, which can cause severe and sometimes fatal envenoming, has been identified as medically relevant already in antiquity, earning scorpions a prominent role in myth and folklore around the globe (Beck and Komposch, 2024; Frembgen, 2004). Moreover, it has been estimated that one third of the global human population is at risk of receiving a scorpion sting and, indeed, several species are recognized as a medical burden in the global south (Chippaux and Goyffon, 2008). Hence, scorpion venom is often perceived as a dangerous and dreadful biomolecular innovation. On the other hand, scorpion toxins are well-known to modulate several biomolecular targets involved in the symptoms and manifestation of diseases (El-Qassas et al., 2024; Ghosh et al., 2019; Suhas, 2022; Uzair et al., 2018). Thus, these toxins are likewise considered a promising source of potential drug leads. Accordingly, an in-depth understanding of the chemical arsenal within scorpion venoms serves a multi-pronged purpose by i) fostering the biological understanding of scorpions, ii) increasing the ability to understand and treat scorpion venom induced pathobiochemistry, and iii) delivering novel valuable bioresources. Modern venomics technologies, particularly venom gland transcriptomics, have recently been optimized to study venom cocktails from hitherto under- or unstudied small arthropods including scorpions (Lüddecke et al., 2019; von Reumont, 2018; von Reumont et al., 2014). For the latter particularly, this has been complemented by the establishment of online resources and data repositories which enable a substantial improvement in the characterization of scorpion venom toxins (Baradaran et al., 2024; Kuzmenkov et al., 2016; Srinivasan et al., 2002). Nevertheless, only a small fraction of global scorpion biodiversity has yet been studied for its venom and the vast majority of scorpion-derived venom toxins remain unknown.

One of the understudied scorpion species is the Black Judaicus scorpion, *Hottentotta judaicus* (Simon, 1872, Figure 1). *Hottentotta judaicus* was originally described as *Buthus judaicus* but later transferred to first *Hottentotta* and then to *Buthotus*. However, the latter placement was made without a clear justification and therefore, Francke (1985) (Francke, 1985) regarded *Buthotus* as a junior synonym of *Hottentotta* (Birula, 1908), leading to the currently accepted combination, *H. judaicus*, although numerous publications still report results from this species under the now outdated *Buthotus judaicus*. Type specimens of *H. judaicus* were originally collected from Jerusalem, the Jordan Valley, and the shores of the Dead Sea. However, this species is known to occur widely across the Middle East, including Jordan, Syria, Lebanon, and Palestine. *H. judaicus* is considered a mesic species, typically inhabiting regions with an annual rainfall of at least 350–400 mm. In its distribution range, this species is commonly encountered, yet its medical significance has not been clarified. While members of *Hottentotta* are often considered dangerous and can potentially cause fatal envenomation, less dramatic symptoms and relatively high LD_50_ values are reported for *H. judaicus* making this species an interesting candidate for investigating venom composition and the evolution of toxin diversity within the genus (Amitai et al., 1981; Basis et al., 2005; West and Hendrickson, 2011).

**Figure 1.**
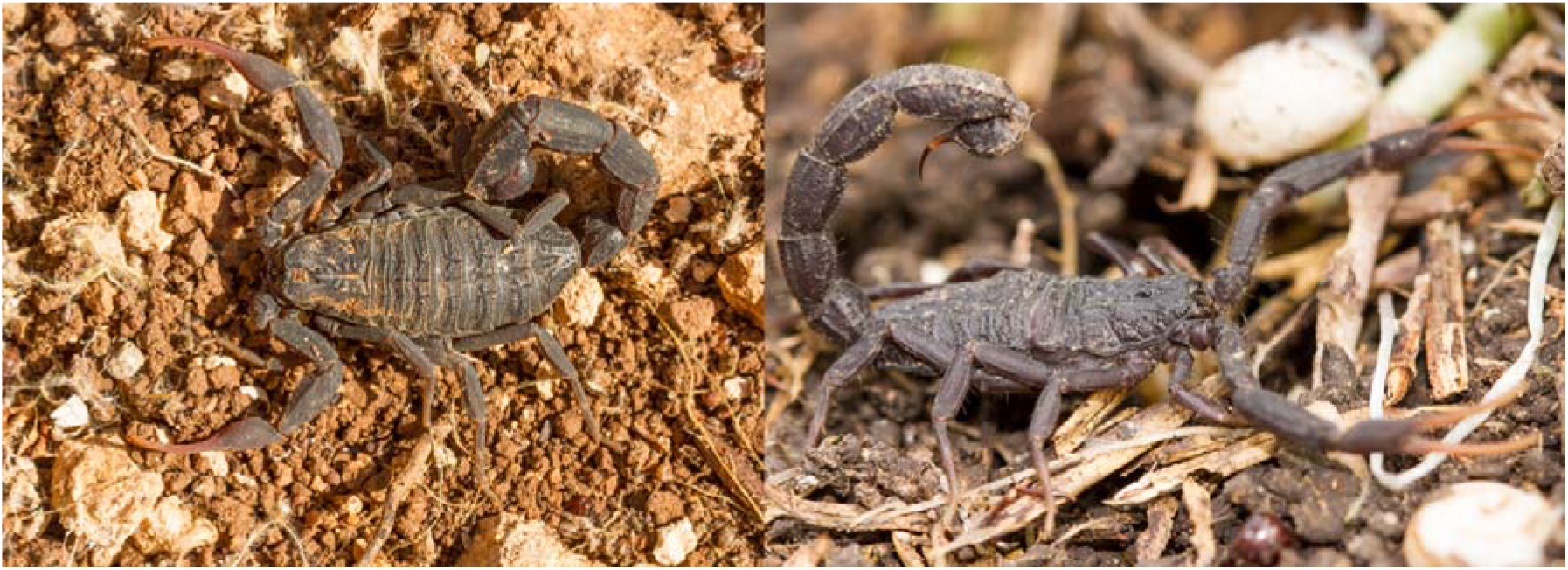
Photographs of adult Black Judaicus scorpions (*Hottentotta judaicus*) in their natural habitat in Israel. Left: dorsal view, right: lateral view. Picture courtesy of: Walter Neser.

That said, this species received relatively little attention for its venom and only few of its toxins have so far been identified (see Table 1). A previous study has investigated its venom profile via sequencing cDNA libraries from a non-replenished venom gland, but an analysis via modern state of the art venomics approaches is currently been missing for this species (Morgenstern et al., 2011).

**Table 1.** Known toxin diversity of *H. judaicus* deposited to the VenomZone database. Given are all entries retrieved for toxins from *H. judaicus* with their respective Uniprot IDs, toxin names, sequence length, and the toxin family they are currently assigned to. For toxins marked with an asterisk (*), only mature sequences are provided while for the others complete precursors including pre- and/or propeptides are listed.

| Uniprot ID | Toxin | Length (#aa) | Toxin family |
| --- | --- | --- | --- |
| F1CIZ6 | Phi-buthitoxin-Hj1a | 64 | Scorpion calcine-like |
| P0DQN8 | Delta-buthitoxin-Hj1a | 66* | Long scorpion toxin (4 C-C) |
| P0DQN9 | Delta-buthitoxin-Hj2a | 64* | Long scorpion toxin (4 C-C) |
| Q56TT9 | Alpha-insect toxin BjaIT | 85 | Long scorpion toxin (4 C-C) |
| P24336 | Beta-insect depressant toxin BjIT2 | 85 | Long scorpion toxin (4 C-C) |
| P56637 | Beta-insect excitatory toxin Bj-xtrIT | 94 | Long scorpion toxin (4 C-C) |
| F1CIY9 | U<sub>10</sub>-buthitoxin-Hj1a | 62 | Short scorpion toxin |
| F1CJ67 | U<sub>10</sub>-hottentoxin-Hj2a | 62 | Short scorpion toxin |
| F1CJ80 | U<sub>10</sub>-hottentoxin-Hj3a | 62 | Short scorpion toxin |
| P83405 | Potassium channel toxin alpha-KTx 9.6 | 28* | Short scorpion toxin |
| P83406 | Potassium channel toxin alpha-KTx 9.7 | 28* | Short scorpion toxin |
| F1CJ89 | Amphipathic peptide Hj0164 | 45 | Non-disulfide-bridged peptide |

We therefore sequenced and analysed the venom gland transcriptome of *H. judaicus* using Illumina technology and a modern bioinformatics pipeline that utilizes multiple assembling algorithms. Our analysis reveals a diverse landscape of toxin-encoding peptides and proteins and shows the presence of several toxins resembling potent homologues from other medically significant scorpions alongside various translationally interesting components. Our exploratory analysis features an important first step towards a more holistic understanding of this species toxin arsenal and facilitates future functional studies of the identified toxins following their biotechnological production.

## 2. Material and Methods

### 2.1 Tissue collection

An adult male specimen of *H. judaicus* was commercially sourced from the pet trade and its identity was morphologically verified. The specimen was kept at ca. 25°C temperature with a 12-h photoperiod for one week in an appropriately sized plastic enclosure (15×15×10 cm) on desert sand with a small cork bark to hide. The scorpion was fed an adult cricket (*Acheta domesticus*), which was subdued via venom injection, to trigger venom expenditure and stimulate the transcriptomic machinery involved in venom replenishment. After 72h the scorpion was anesthetized with CO_2_ and the venom glands were dissected from its telson under a Stemi 508 stereomicroscope (Zeiss), washed in distilled water, and transferred to 1 ml RNAlater solution (Sigma-Aldrich). The sample was stored at 4°C until further processing.

### 2.2 Transcriptome sequencing

RNA extraction and sequencing were outsourced to Macrogen. Following RNA extraction, libraries were constructed using the TruSeq Stranded mRNA Library Prep Kit (paired-end, 151-bp read length) (Illumina). Quality was controlled by the verification of PCR-enriched fragment sizes using an Agilent Technologies 2100 Bioanalyzer with a DNA 1000 chip. The library quantity was determined by qPCR using the rapid library standard quantification solution and calculator (Roche). The libraries were sequenced using Illumina technologies, further details are provided in in Supplementary Table S1. Raw sequencing data are available at the European Nucleotide Archive (ENA) (Study PRJEB90058).

### 2.3 Transcriptome assembly

Transcriptome data were processed using a modified version of our in-house assembly and annotation pipeline (Hurka et al., 2022). All input sequences were inspected using FastQC v0.12.1 (https://www.bioinformatics.babraham.ac.uk/projects/fastqc/) before trimming with cutadapt v4.9 (Martin, 2011) using a quality cutoff of 28 and a minimum length of 25 bp. The trimmed reads were corrected using Rcorrector v1.0.7 (Song and Florea, 2015) and assembled *de novo* using a pipeline incorporating Trinity v2.15.1 (Haas et al., 2013) with a minimum contig size of 30 bp and maximum read normalization of 50 and rnaSPAdes v3.15.5 (Bushmanova et al., 2019) with and without error corrected reads. Both rnaSPAdes assemblies were built with k-mer sizes of 21, 33 and 55. All contigs were combined into a single assembly with identical transcripts from all assemblers being merged with fastanrdb v2.4.0 and strand specificity was taken into account (Slater and Birney, 2005). The reads were remapped to the assembly using HISAT2 v2.2.1 (Kim et al., 2019) and expression values (transcripts per million, TPM) were calculated using StringTie v2.2.2 (Kovaka et al., 2019). SAM and BAM files were converted using samtools v1.20 (Danecek et al., 2021). Open reading frames (ORFs) were then predicted with TransDecoder v5.7.1 (Haas et al., 2013) with a minimum length of 10 amino acids.

### 2.4 Annotation

Identified toxin precursors were annotated using InterProScan v5.69-101.0 (Jones et al., 2014) and a DIAMOND v2.1.9 blastp search (Buchfink et al., 2021) against the public available databases VenomZone (“VenomZone,” 2025), UniProtKB/Swiss-Prot Tox-Prot (Jungo and Bairoch, 2005), UniProtKB/Swiss-Prot and UniProtKB/TrEMBL v2024_04 (The UniProt Consortium, 2019) was performed. The E-value was set to a maximum of 1 × 103 in ultra-sensitive mode with all target sequences reported (--max-target-seqs 0). We then calculated the coverage of query and subject, and the similarity with the BLOSUM62 matrix using BioPython v1.83 (Cock et al., 2009) for each hit. Sorting by similarity, query and subject coverage for each toxin candidate led to the resulting hit for the final analysis. Precursors without a predicted signal peptide by SignalP v6.0h (Teufel et al., 2022) in slow–sequential mode for eukarya were removed from our dataset. Annotated precursors were aligned to known venom components from the same putative family to verify our assignments using ClustalW in Geneious v10.2.6 (www.geneious.com). The retrieved bona fide peptides were provided with an identifier based on the species name (Hju = *Hottentotta judaicus*) combined with a unique number. The resulting annotated peptide sequences are provided in Supplementary Table S1.

## 3. Results and Discussion

### 3.1 High quality H. judaicus venom gland transcriptome library

The generated paired-end libraries were checked for DNA quality and quantity prior further processing. The yielded library contained a total of 42,910,214 paired-end reads, characterized by a GC content of 40.2%, a Q20 of 98.2% and Q30 of 95.3% respectively. These data indicated that a high-quality transcriptome library was generated, therefore it was subsequently sequenced and analyzed via our custom-made pipeline.

### 3.2 The venom profile of H. judaicus revealed by transcriptomics

After assembling the *H. judaicus* venom gland transcriptome, we annotated the generated sequences via multiple approaches, including BLAST searches, InterProScan, and comparative alignments, to identify bona fide venom components. A qualitative assessment of the *H. judaicus* venom gland transcriptome was then calculated. Therefore, we calculated the total number of identified different transcripts. Then, we counted the total number of transcripts within each toxin family based upon our sequence annotation results. The percentage of diversity within each toxin family was expressed as the relative percentage of all transcripts within a family divided by the total number of different transcripts identified. Remapping of reads to the gathered assembly further allowed us to calculate the abundance of transcripts as transcripts per million (TPM). Summed up TPMs within each putative protein family, divided by the total summed up TPM of all venom components, were used to gather a quantitative perspective on the venom profile. In total, we identified 305 transcripts encoding potential venom components that were assigned to 19 distinct putative protein families.

First, we analyzed the retrieved venom profile in terms of encoded diversity of protein families (Figure 2A). The most diverse group, containing 21.3% of all transcripts were enzymes and other high molecular weight proteins, followed by short and long scorpion toxins (4 C-C type) that each contained 19.7% of all transcripts, respectively. The next diverse protein category was an assembly of various minor scorpion venom protein families referred to as “other”. These together represent 18.4% of all transcripts and included TIL- and Kunitz-type serine protease inhibitors, scorpion calcin-like, and venom protein 30.1-like proteins among others. Potential antimicrobial peptides (AMPs), including linear and disulfide crosslinked types, featured 11.8% of all transcripts, while long scorpion toxins (3 C-C type) and long chain scorpion toxins featured 7.9% and 1.3% respectively. On the level of abundance, we identified the short scorpion toxin family as the most prevalent group of venom components that constitute 37.2% of TPM (Figure 2B). Second to these were long scorpion toxins (4 C-C type) with 23.1% followed by enzymes plus other high molecular weight proteins with 19.4% of the total TPMs. Long scorpion toxins (3 C-C type) featured 13.8% of the TPMs, while putative AMPs accounted for 3.1% of the transcript abundances. A marginal 0.6% of the TPM were assigned to long chain scorpion toxins and a total of 2.9% of all TPMs were assigned to others.

**Figure 2.**
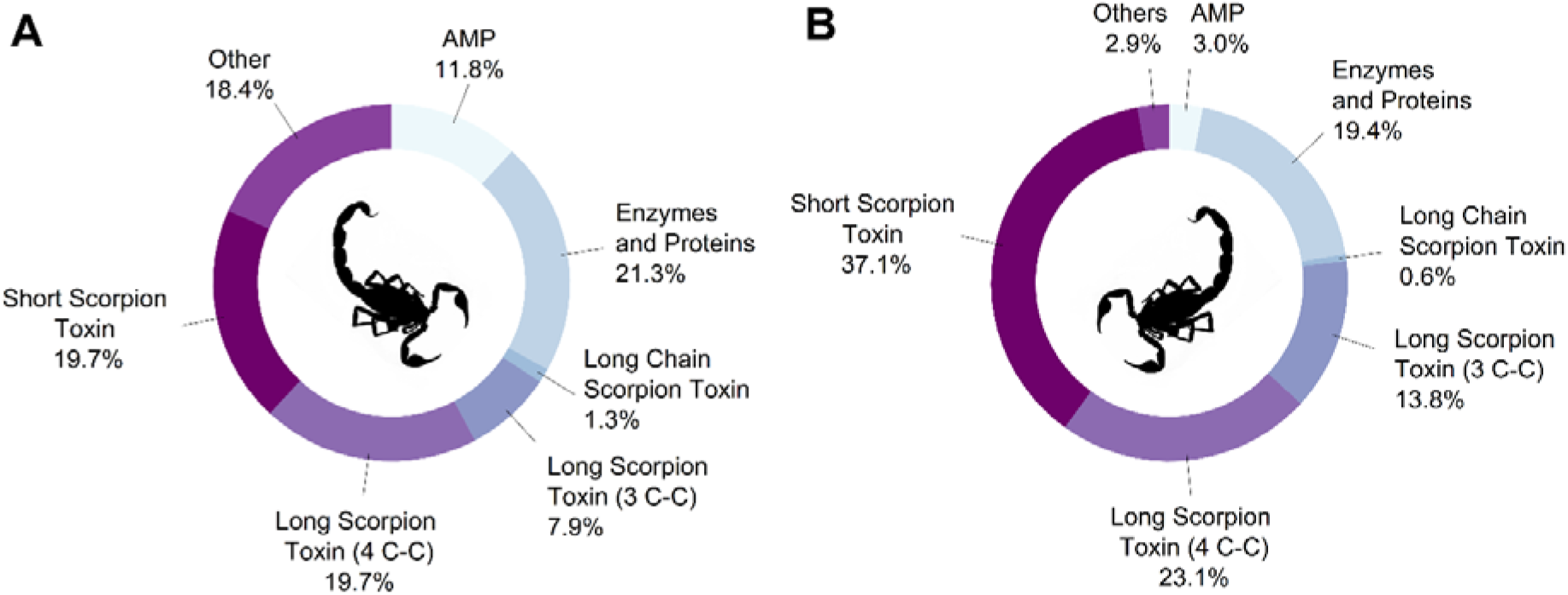
The toxin arsenal of *H. judaicus* inferred by transcriptomics. The *in silico* determined profile of toxin encoding precursors is illustrated on the level of diversity (A, based on number of precursors per family relative to all precursors of the transcriptome) and abundance (B, percentage of summed up TPM of precursors within each family relative to all TPM). Scorpion modified after Gareth Monger©.

### 3.3 Major toxin families and their activities

The venom gland transcriptome of *H. judaicus* abounds with a high degree of transcript diversity and contains mRNAs encoding for hundreds of compounds. Four groups of components dominate the venom profile of this species: short scorpion toxins, long scorpion toxins of 3 C-C and 4 C-C type, as well as enzymes and other proteins of higher molecular weight. Their potential activities and supposed biological role of the encoded *H. judaicus* toxins are discussed henceforth.

### 3.3.1 Short scorpion toxins

The short scorpion toxin superfamily is one of the most diverse families of scorpion toxins and very abundant in venoms from the Buthidae family (Di Nicola et al., 2024). They are relatively short neurotoxic peptides with mature proteins usually ranging between 30-40 amino acids (Dardevet et al., 2015; Saucedo et al., 2012). Structurally, these toxins typically contain an alpha-helix and an antiparallel beta-sheet which are covalently linked through three disulfide bonds. This structure is commonly referred to as CS-alpha/beta fold (Saucedo et al., 2012). Short scorpion toxins mostly block various types of potassium channels, including voltage-gated-, and calcium-activated potassium channels (Saucedo et al., 2012). Through the inhibition of targeted potassium channels, these toxins block the signal transduction within the central nervous system to cause paralysis and thereby facilitate fast prey inactivation. Furthermore, several short scorpion toxins are known to interact with chloride and calcium channels, though the exact mechanism of interaction requires further investigation. These chloride channel interacting toxins, including its most prominent member Chlorotoxin from the deathstalker *Leiurus quinquestriatus* (Ehrenberg, 1828), are of great translational interest as anticancer agents, yet their biological role remains to be resolved (Cohen-Inbar and Zaaroor, 2016; Ojeda et al., 2016).

Several transcripts have similarities with pharmacologically interesting scorpion toxins (Figure 3). For instance, transcripts Hju186 and Hju194 have sequence similarity to potassium channel toxin alpha-KTx 15.3 from highly toxic Moroccan fat-tailed scorpion (*Androctonus mauritanicus* [Pocock, 1902]) venom. This toxin is an inhibitor of A-type voltage-gated potassium channels of striated neurons but also interferes with ERG1/Kv11.1/KCNH2 potassium channels (Maffie et al., 2013; Zoukimian et al., 2019). Other transcripts (e.g. Hju357, Hju358, and Hju359) exhibit sequence similarity to potassium channel toxin alpha-KTx 8.5 from the yellow Iranian scorpion (*Odonthobutus doriae* [Thorell, 1876]). This toxin is also a selective inhibitor of a voltage-gated potassium channel (Kv1.2/KCNA2) (Abdel-Mottaleb et al., 2006). Transcript Hju345 shares similarity with potassium channel toxin alpha-KTx 1.1 from the medically relevant Hebrew deathstalker scorpion (*Leiurus hebraeus* [Birula, 1908]). Interestingly, this toxin inhibits a wide range of potassium channels at nanomolar ranges, including shaker, Kv1.2/KCNA2, Kv1.3/KCNA3 and several others (Kasheverov et al., 2019). It also has weak inhibitory effects on nicotinic acetylcholine receptors and antimicrobial properties (Kasheverov et al., 2019; Yount and Yeaman, 2004).

**Figure 3:**
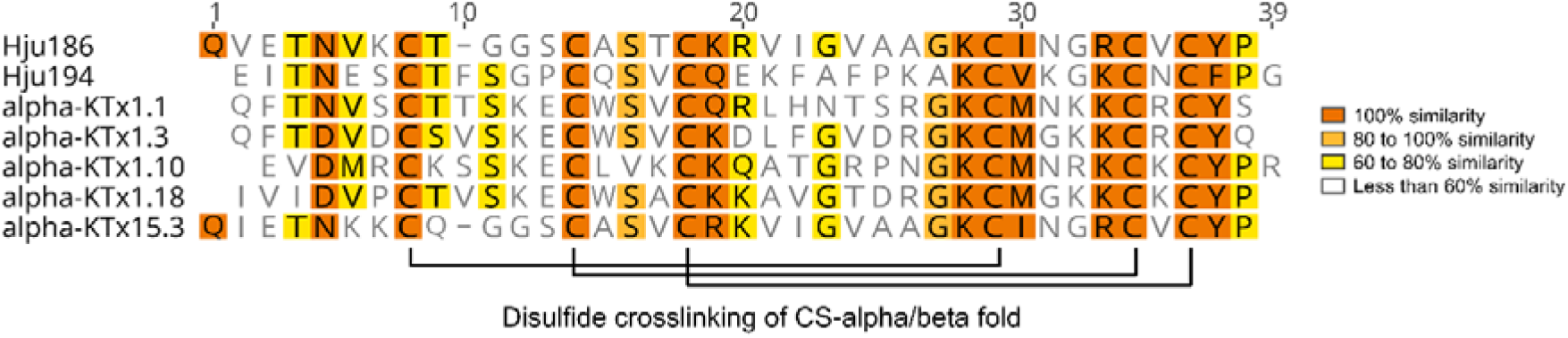
Alignments of short scorpion toxins. Selected sequences identified in this study aligned to known members of the family including the corresponding disulfide crosslinking of the CS-alpha/beta fold. The color code indicates sequence similarity per position in the alignment.

Lastly, several transcripts have similarity to short scorpion toxins previously described from *Hottentotta* venom. Some transcripts resemble *H. judaicus* toxins, including Hju411, Hju416, Hju417, and Hju418, which are identical or similar to U10-buthitoxin-Hj1a, an uncharacterized short scorpion toxin, as well as the transcripts Hju199, Hju353, Hju354, Hju355, and Hju360, which has similarity to either potassium channel toxin alpha-KTx 9.6, an activator of calcium channels and ryanodine receptors (Zhu et al., 2004). That said, some transcripts display similarity to toxins from the Indian red scorpion (*H. tamulus* [Fabricius, 1798]). This includes transcript Hju111, which mirrors Potassium channel toxin alpha-KTx 5.5, a blocker of small conductance calcium-activated potassium channels (Pedarzani et al., 2002). Transcripts Hju324, Hju325, and Hju326 feature similarity to lepidopteran-selective toxin CTXL, a neurotoxin with chloride channel modulatory activities in glioma cells and high toxicity towards *Chloroidea virescens* (Fabricius, 1777) caterpillars, but not against dipteran and mouse models (Wudayagiri et al., 2001). Some other transcripts, including, Hju177 and Hju330 display similarity to neurotoxin BtITx3, a toxin with similar pharmacological activity as CTXL when tested in glioma cells, but with severe neurotoxicity in *Helicoverpa armigera* (Hübner, 1809) caterpillars (Dhawan et al., 2002).

### 3.3.2 Long scorpion toxins

Long scorpion toxins fall into two distinct categories, the 3 C-C and the 4 C-C type depending on the number of disulfide bonds present in each peptide (Saucedo et al., 2012). Both types are primarily targeting sodium channels albeit particularly 3 C-C-type toxins are known to further modulate potassium channel (Saucedo et al., 2012). Based upon their neurotoxic activities, these toxins are mainly used for prey subjugation. Both types of long scorpion toxins feature major components of the *H. judaicus* venom gland transcriptome, and exhibit similarity with known potent homologues from other scorpions.

For the 3 C-C type, this includes several transcripts. For instance, transcripts Hju83, Hju84, Hju94 and others are quite similar to the toxin Ts28 from Brazilian scorpion (*Tityus serrulatus*, Lutz & Mello, 1922) venom (Figure 4A) which is thought to modulate voltage-gated sodium channels and to cause lipolysis in rat adipocytes following the formation of a heterodimer (Kalapothakis et al., 2021). Several transcripts within our transcriptome show sequence similarity to this peptide, including Hju123, Hju131, and Hju219. Finally, transcripts Hju172, Hju173, and Hju320, plus others, show sequence similarity to toxin Acra III-2 from the Arabian fat-tailed scorpion *Androctonus crassicauda* (Olivier, 1807), a potential modulator of voltage-gated sodium channels.

**Figure 4:**
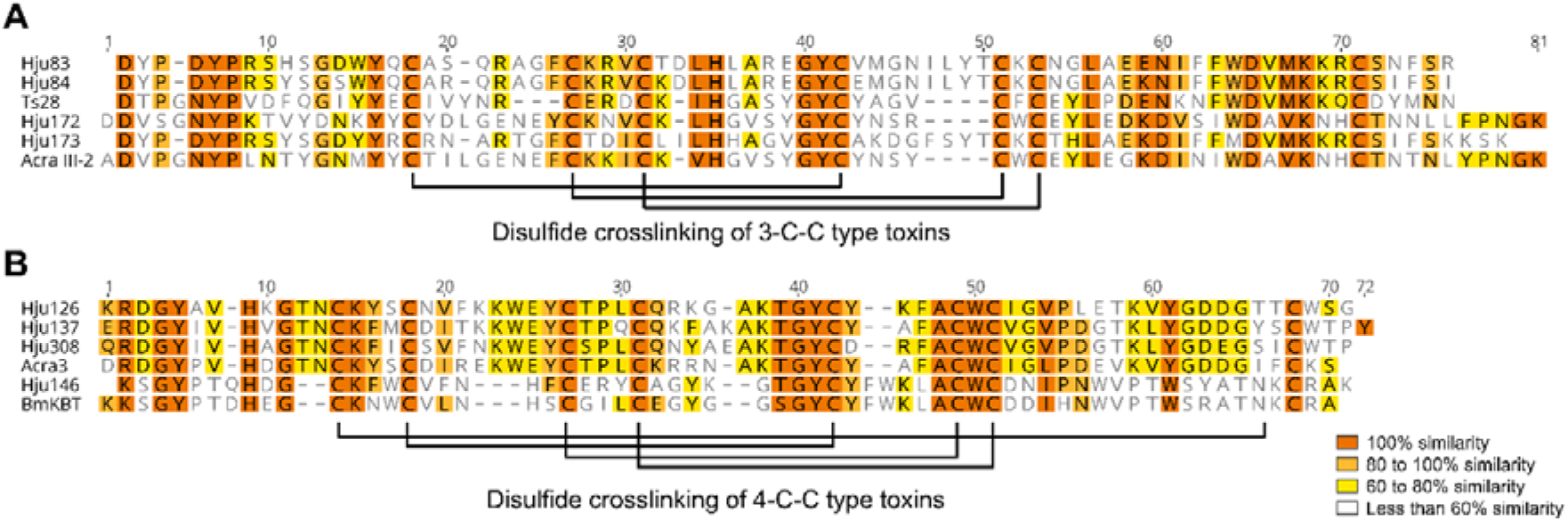
Alignments of long scorpion toxins. Selected sequences (A: 3 C-C type toxins; B: 4 C-C type toxins) identified in this study aligned to known members of the family. The color code indicates sequence similarity per position in the alignment.

A noteworthy degree of transcripts was assigned to 4 C-C long scorpion toxins. Several resemble the toxin Acra3 from *A. crassicauda* (e.g. Hju126, Hju137, or Hju308, Figure 4B). This toxin is known to cause severe, often fatal, excitatory and convulsive neurotoxicity in mice *in vivo* and also to exert potent cytotoxicity in BC3H1 mouse brain tumor cells (Caliskan et al., 2013, 2012). Furthermore, several transcripts (e.g. Hju136, Hju309, and Hju222) have similarity to the uncharacterized toxin Acra5, also stemming from *A. crassicauda*, which based on its sequence similarity is presumed to be a potential inhibitor of activation for voltage gated sodium channels and thus may block neuronal transmission. A similar mode of action might be exerted by MeuNaTxalpha-3 from Lesser Asian scorpion (*Mesobuthus eupeus* [C.L. Koch, 1839]) which is similar to some of the *H. judaicus* transcripts, particularly to Hju148, Hju174, and Hju307. A noteworthy degree of transcripts, such as Hju146, or Hju147, shows sequence similarity to BmKBT from the Manchurian scorpion *Olivierus martensi* [Karsch, 1879]. This toxin is a putative sodium channel inhibitor causing severe toxicity in rodents but non-lethal paralysis in lepidopteran insects (Ji et al., 2002). Further, the transcripts Hju213, Hju227 and Hju306, have similarity to beta-mammal/insect toxin Lqhb1 from Hebrew deathstalker scorpion (*L. hebraeus*). This toxin interacts with voltage-gated sodium channels by shifting the voltage of activation towards more negative potentials and perturbs the channels activation, leading to uncontrolled, repetitive signal release (Gordon et al., 2003). Lastly, several transcripts show similarity to insect toxin BsIT3 from the Sind red scorpion (*Hottentotta tamulus sindicus* [Fabricius, 1798]), e.g. Hju157, Hju158, and Hju159. This toxin is a depressant toxin that first facilitates transient contractile paralysis followed by flaccid paralysis via interaction with voltage-gated sodium channels where they shift the voltage of activation (Ali et al., 2001). These toxins are only active in insects (Ali et al., 2001).

### 3.3.3 Enzymes and other proteins

Enzymes and other proteins make up the most diverse and third most abundant category found in the venom gland transcriptome. Transcripts belong to several enzyme families, such as chitinases, hyaluronidases, phospholipase A_2_, as well as members of the M12 metalloprotease family (M12A/astacin, M12B/reprolysin). Besides them, proteins from the CAP superfamily (Cysteine-rich secretory proteins, Antigen 5 and Pathogenesis-related protein 1) are present. A proteomic analysis of venom from six scorpion families has revealed the presence of an enzymatic core consisting of 24 distinct enzyme families found in all examined scorpion venoms. With the exception of CAP and reprolysin, all remaining enzymes identified in our transcriptome are part of this enzymatic core and have been assigned to potential functions and biological roles (Delgado-Prudencio et al., 2022).

Chitinases are minor components of the *H. judaicus* venom gland transcriptome (1.3% of transcript diversity and 0.1% of summed-up TMP). Following the proposed classification system, chitinases degrade the chitin exoskeletons of arthropods and may aid in the liquefication of prey tissue before ingestion (Fuzita et al., 2015). Hyaluronidases account for 1.3% of all transcripts and 1.4% of TPM respectively and may facilitate the diffusion of venom components (Delgado-Prudencio et al., 2022). They hydrolyze hyaluronic acid are therefore presumed to act as spreading factors (Ferrer et al., 2013). However, this process is restricted to vertebrates, as the extracellular matrix of insects lacks hyaluronic acid. Therefore, they may be utilized defensively against vertebrate predators (Dresler et al., 2024a). Phospholipases A_2_ (PLA_2_) represent 3.3% of all transcripts and 0.2% of all TPM. They are of small molecular size (< 20 kDa) and common venom components in its secretory form, sPLA_2_ (Dresler et al., 2024a). They catalyze the hydrolysis of membrane glycerophospholipids at the sn-2 position causing the release of fatty acids and lysophospholipids (Murakami and Kudo, 2002). Structural analysis has shown that all scorpion venom PLA_2_ are calcium dependent (Krayem and Gargouri, 2020). Scorpion PLA_2_ are considered to be pre-digestive enzymes and spreading factors(Delgado-Prudencio et al., 2022). Metalloproteases, such as astacins and reprolysin, account for 15.1% of all identified transcripts and 10.5% of summed up TPM. They degrade extra cellular matrix components and may act as spreading factors or pre-digestive enzymes (Dresler et al., 2024a; Medina-Santos et al., 2019).

Besides components known for their enzymatic activity, CAP proteins are still awaiting clarification of their potential enzymatic activities. They account for 3.9% of all transcripts and 7.2% of the TPMs and are generally not considered as enzymatic, but some CAP proteins described in spider venom show sequence similarities to proteolytic cone snail CAPs (Dresler et al., 2024a; Lüddecke et al., 2020). Based on that similarity, they have been speculated to exert proteolytic activity, but functional characterizations are still pending (Lüddecke et al., 2020). Scorpion CAP proteins display sequence similarity to spider venom CAPs and therefore might cause similar reactions (Dresler et al., 2024a).

### 3.4 The translational potential of Hottentotta judaicus venom

The analysis of the *H. judaicus* venom gland transcriptome revealed a seemingly complex venom profile containing a large proportion of neurotoxins and enzymes. Several of the identified components may offer translational avenues. The areas of particular interest revolve around biomedicine, agriculture, as well as industrial production and are discussed henceforth.

Scorpion toxins have emerged as an attractive starting point to identify drug leads (Ortiz et al., 2015; Uzair et al., 2018; Xia et al., 2023). The herein presented venom gland transcriptome abounds with a high degree of putative neurotoxins assigned to pharmacologically interesting toxin families that are worthy further investigation. Leveraging biotechnological production and accessing these molecules on laboratory-scale will yield may yield novel hit components with desired pharmacological profiles (Lüddecke et al., 2023b). These may be further designed and developed towards pharmaceutical utilization. That said, besides the neurotoxins we identified a range of linear toxins that represent potential AMPs. Such AMPs, particularly from arachnid venom, are known source of anti-infective peptides that target a wide range of pathogenic bacteria and other microbes (Almaaytah and Albalas, 2014; Lüddecke et al., 2023a; Saez et al., 2010; Xia et al., 2024). Several arachnid venom AMPs are known to potently target even multi-drug resistant bacteria (Erkoc et al., 2024) and our analysis reveals that *H. judaicus* features a so far neglected source of anti-infective leads.

Another area of utilization are agricultural applications (King, 2019; King and Hardy, 2013; Saez and Herzig, 2019). Over the course of evolution, arachnid venoms have been refined to target insect receptors (Windley et al., 2012). Paired with their biodegradability, they are rendered excellent templates for the creation of eco-friendly bioinsecticides (Herzig et al., 2025; Windley et al., 2012). In that context, it appears that *H. judaicus* produces agriculturally interesting toxins, as several of the identified transcripts appear to be insect-selective neurotoxins that could be explored further (such as the transcripts with similarity to insect selective toxins CTXL, BtITx3, and BsIT3 as outlined in section 3.3.1 and 3.3.2).

Moreover, previous studies suggested, that scorpion venom AMPs can be used to kill pest insects that depend on functional gut microbiomes (Luna-Ramirez et al., 2017). Facing the diversity of AMP-like toxins identified in our venom gland transcriptome, it would be a promising effort to screen them for targeting microbiome-depending pest insects to explore their agricultural potential.

Lastly, we have identified a range of enzymatic components in the *H. judaicus* transcriptome. Their presence in venoms from scorpions and other arachnids has been proposed as one of the grand challenges in arachnid toxinology (Delgado-Prudencio et al., 2022; Dresler et al., 2024a, 2024b; Herzig, 2023). Enzymes are important elements for several bioeconomic areas, particularly for the production of industrial goods (Mesbah, 2022; Singh et al., 2016). Given their efficiency, sustainability and ability to catalyze challenging chemical reactions, novel enzyme-technologies are constantly sought after in various industries (Mesbah, 2022; Singh et al., 2016). With that in mind, the components identified in *H. judaicus*, are worthy of further investigations, which could lead to novel production tools.

### 3.6 Challenges and future perspectives

While the diversity of potentially novel toxins with similarity to translationally attractive homologues suggest a bioeconomic potential, our data needs to be interpreted with some caution. First, our analysis is based solely on transcriptomic evidence and not all sequenced precursors may be present on proteome level. It has recently been shown that transcriptomic data alone overestimates the true diversity of a venom profile and may create false-positive precursors (Smith and Undheim, 2018). It will therefore be important to consolidate the presence of potentially interesting transcripts on the protein level by leveraging mass spectrometry technologies. A second hurdle is presented by the displayed degree of similarity. Whilst all identified toxins appear to be members of well-characterized protein families, this does not necessarily allow direct conclusions on their bioactivity and avenues of utilization. Some of the herein discussed toxins have similarity to toxins previously classified as translationally attractive, yet albeit these usually emerged as the most similar known peptides during BLAST searches (see Supplementary Table S1), their overall similarity is not always very high (Table 2). Particularly with venom peptides it has been repeatedly shown that single amino acid exchanges can already have tremendous functional repercussions and can alter the mode of action, selectivity, and potency of toxins (Panchera et al., 2025; Sunagar et al., 2013; Undheim et al., 2015; Yin et al., 2020). Hence, it will be of utmost importance to evaluate the herein identified toxins of potential bioeconomic value based on experimental screening before any a priori assumptions can be made.

**Table 2.**
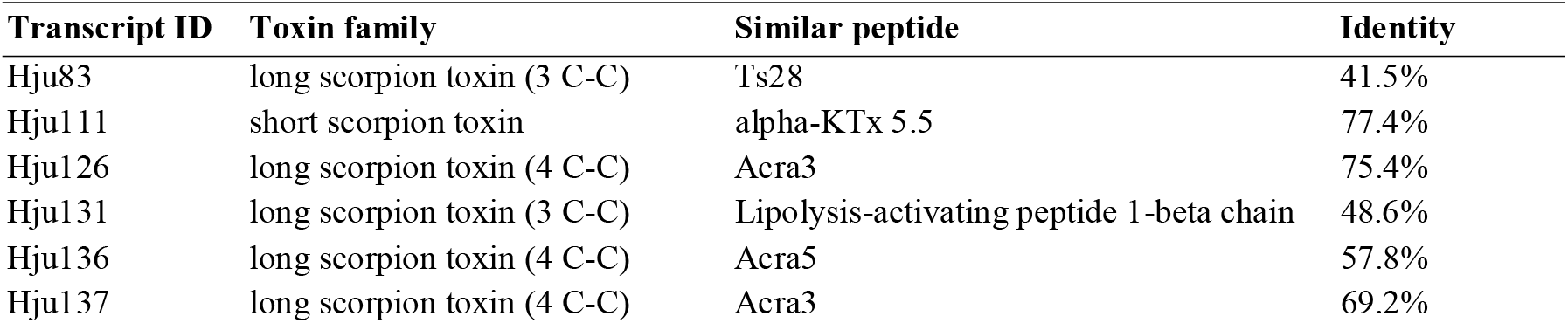
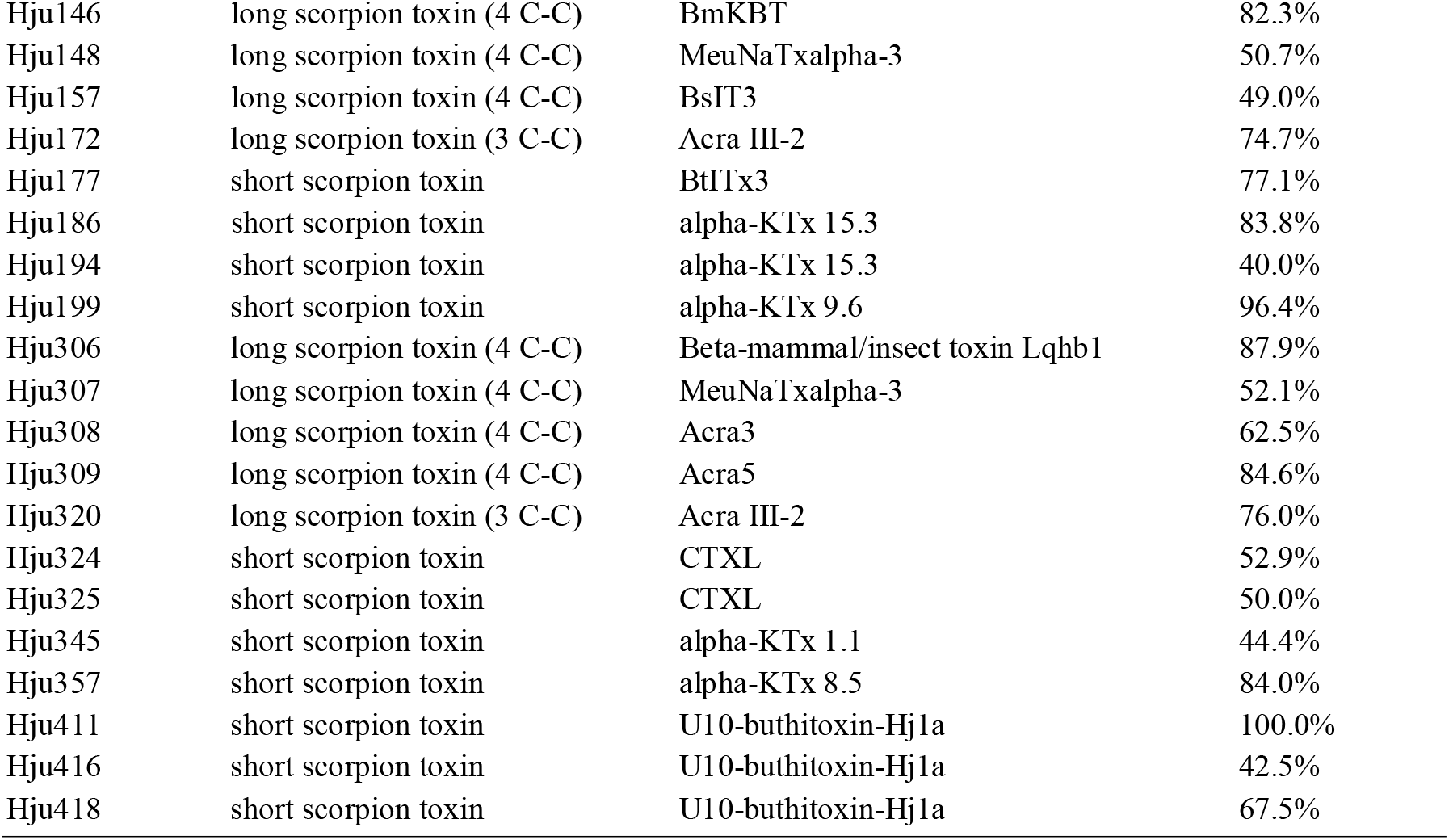
Similarity of selected noteworthy transcripts to known scorpion venom toxins. The table lists transcript IDs with their respective toxin family, the most similar known peptide and their degree of identity on amino acid level.

## 4. Conclusion

Here, we present the venom gland transcriptome of *H. judaicus* one of the still neglected scorpions from the Middle East. We show that its venom contains a large library of toxins resembling highly toxic peptides from closely related taxa, which represents an important initial step to understand the pathobiochemistry of this species and to unveil the most important toxins for future in-depth examination. Also, we show that several toxins appear as attractive study objects for translational projects revolving around drug discovery, agriculture, and industrial production. We recommend that future studies employ venom proteomics to verify the existence of the most promising components within the venom of *H. judaicus and* aim at producing these toxins using biotechnological approaches and explore their functional space to untap these promising bioresources for the bioeconomy.

## Acknowledgements

We acknowledge technical assistance of the Bioinformatics Core Facility at the professorship of Systems Biology at JLU Giessen and the provision of IT resources and general support by de.NBI/ELIXIR-DE (W-de.NBI-010) funded by the Federal Ministry of Education and Research.

## Appendix A

Supplementary data

Supplementary data to this article can be found online at PRJEB90058.

